# Water body size shapes body size evolution via subgenomic selection in allopolyploid fish

**DOI:** 10.1101/2025.09.22.677753

**Authors:** Li Ren, Yiyan Zeng, He Dai, Yakui Tai, Kaikun Luo, Shaofang He, Xin Gao, Jialin Cui, Mengxue Luo, Xueyin Zhang, Hong Zhang, Mengdan Li, Ling Liu, Hailu Zhou, Jinhui Zhang, Xiaohuan Han, Shi Wang, Shaojun Liu

## Abstract

Allopolyploidization is a major force in species evolution, and studying allopolyploids in diverse environments provides insights into their genetic mechanisms. Here, we investigated how water body size variation influences body size in allopolyploid fish. We used a nascent allopolyploid lineage derived from goldfish (*Carassius auratus*) and common carp (*Cyprinus carpio*) and established two populations in small and large water bodies for two generations. Genomic, transcriptomic, and epigenomic analyses revealed that the larger population had higher genetic diversity in subgenome R, while the smaller population had higher diversity in subgenome C. Potential selection regions were associated with growth regulatory genes, suggesting a role for gene regulation in body size adaptation. Dynamic gene expression patterns in various tissues indicated potential mechanisms underlying muscle development and adaptation to smaller water bodies. Furthermore, selection regions were linked to changes in chromatin accessibility, with subgenome R being more susceptible to these changes during genetic divergence. Our findings demonstrate that water body size variation drives body size in allopolyploid fish through subgenomic selection, impacting genetic variation, gene regulation, and epigenetic mechanisms.

## Introduction

Allopolyploidization, a process that involves the hybridization of two distinct species followed by a doubling of chromosome number, is a significant evolutionary phenomenon that has led to the emergence of new species (Comber & Smith, 2004; Li Ren et al., 2022). This process has been linked to the origins of various species, including gibel carp (*Carassius gibelio*) (Wang et al., 2022), goldfish (*Carassius auratus*) (Luo et al., 2020), and common carp (*Cyprinus carpio*) (P. Xu et al., 2019). They belong to the Cyprininae subfamily, which is recognized as the largest and most diverse family of fish (Yang et al., 2015). The cyprinid fish exhibit remarkable morphological diversity (Nelson, Grande, & Wilson, 2016), including body lengths that range from about 8 mm for *Paedocypris progenetica* to approximately 3 m for *Catlocarpio siamensis* (He et al., 2019; Jacquemin & Pyron, 2016; Kottelat, Britz, Hui, & Witte, 2006). Goldfish and common carp share a common ancestor arising from an allopolyploidization event that occurred approximately 13.75 million years ago (Mya), followed by a subsequent divergence at 9.95 Mya (D. Chen et al., 2020; Luo et al., 2020; P. Xu et al., 2019; P. Xu et al., 2014). Despite sharing numerous biological characteristics, such as omnivorous behavior, vibrant body coloration, and broad temperature adaptability, they exhibit distinct differences in terms of body size and barbel morphology (Luo et al., 2020; P. Xu et al., 2014). This phenomenon offers insights into how the effects of allopolyploidization shape the phenotypic evolution, particularly in terms of body size, within specific environments.

Compared to determinate growth patterns observed in other vertebrates like mammals and birds, fish exhibit indeterminate growth, influenced by a multitude of factors (Amson & Bibi, 2021). These factors include temperature, food availability, light regime, oxygen levels, predator density, intraspecific social interactions, and genetics (Albert & Johnson, 2012; Pinto-Coelho, Martins, & Guimaraes Junior, 2021; Uyeda, Pennell, Miller, Maia, & McClain, 2017). Moreover, the body size of fish is intricately linked to the dimensions of their aquatic environments (Scheffer et al., 2006). This reflects adaptive evolutionary strategies employed by different species in specific environments. Smaller fish species typically employ evolutionary tactics such as rapid reproduction, shorter lifespans, and adaptable feeding habits (Olden, Hogan, & Zanden, 2007). These strategies enable them to confront challenges posed by intense competition, predation pressure, and unpredictable environments. On the other hand, larger fish may adopt more stable survival strategies. These strategies may include slower reproduction, longer lifespans, and specialized feeding behaviors (Olden et al., 2007). Such adaptations empower larger fish to navigate through relatively stable yet resource-limited ecosystems.

Allopolyploids, in comparison to their diploid ancestors, display significant genome plasticity and redundancy (Luo et al., 2020; P. Xu et al., 2019). This plasticity may be associated with active transposable elements, a regulatory network derived from homologous genes, and diverse patterns of DNA methylation regulation (Li Ren et al., 2022). Various genomic changes, including DNA recombination and alterations in epigenetic and gene expression levels, have been reported in some plants, such as Brassica (Chalhoub et al., 2014), rice (N. Li et al., 2019), and cotton (Song, Zhang, Stelly, & Chen, 2017). The diversified changes among subgenomes constitute a pivotal genetic foundation facilitating rapid phenotypic variation in the domestication of polyploid crops (Akagi, Jung, Masuda, & Shimizu, 2022). Despite this, there has been relatively little exploration of the role of allopolyploidization in the evolution of vertebrates.

Nascent allopolyploid fish will be an effective model for studying the effects of allopolyploidization and the evolution of body size. This is because this stage contains the genetic material of the parental species in a more complete form, and its genome plasticity is relatively large. A nascent allopolyploid population (4nR_2_C_2_) was successfully established by the hybridization of female goldfish and male common carp and subsequent whole genome duplication (Shaojun Liu et al., 2001; S. Liu et al., 2016). After multiple generations of self-fertilization, this genetic population showed rich genetic diversity and plasticity (Li Ren et al., 2022). This study is the first to cultivate two allopolyploid populations in environments that differed solely in water body size. All other environmental factors, including illumination, water temperature, fish density, oxygen content, and food supply, were identical. After two generations, we observed that the individual sizes of different populations and within populations showed diversified characteristics under different water environments. To decipher the genetic mechanisms behind this phenomenon, we conducted population genetics analyses based on the whole genome resequencing of 200 individuals. To further explore the mechanism by which water environments regulate gene expression through epigenetic modifications, we also conducted studies on chromatin accessibility in muscle tissue. These studies will provide us with effective approaches to exploring the relationship between water body size and the evolution of fish body size.

## Results

### SNPs and CNVs

To investigate how the nascent allotetraploid fish, 4nR_2_C_2_ (derived from the hybridization of goldfish (*Carassius auratus*) and common carp (*Cyprinus carpio*)), adapts to different water body sizes, we established two populations across two generations (Fig. 1A). These populations (4nR_2_C_2_-B and 4nR_2_C_2_-S) were bred in pools with contrasting water volumes (8125 m³ for 4nR_2_C_2_-B and 400 m³ for 4nR_2_C_2_-S). Importantly, all other environmental conditions were meticulously maintained to be identical for both populations. The F_2_ generation of the 4nR_2_C_2_-B population exhibited a larger body size compared to the 4nR_2_C_2_-S population (Fig. 1B).

**Fig. 1.**
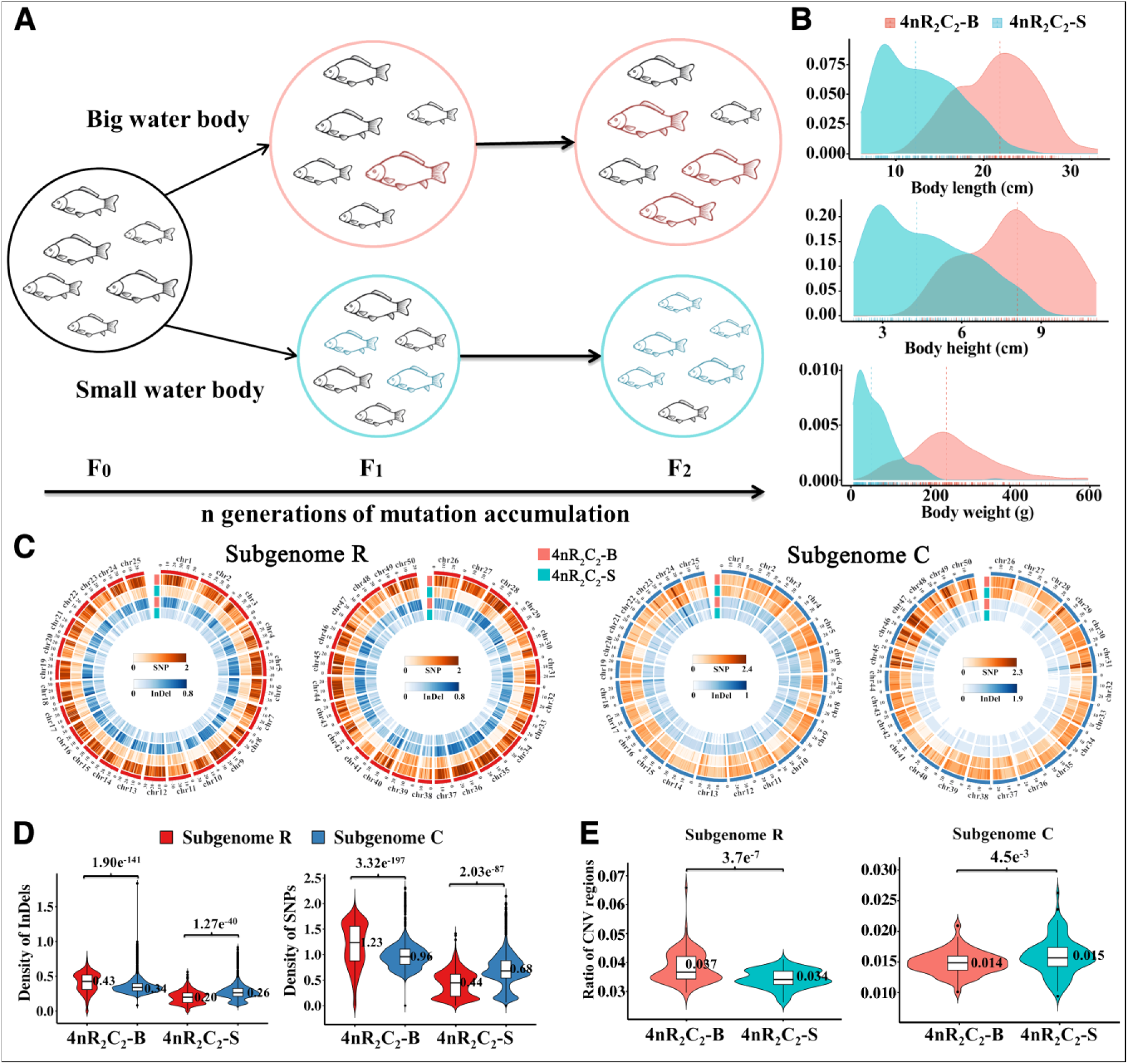
Divergence of nascent allopolyploid lineages in different water body sizes. (**A**) Mutation accumulation in two populations across generations. (**B**) The growth traits of the two populations, including body length, height, and weight, were detected in F_2_ generations (12 months after hatching). (**C**) SNPs and InDels distributions in each chromosome for 4nR_2_C_2_-B and 4nR_2_C_2_-S populations. (**D**) Distribution of InDels and SNPs between subgenomes R and C. A *t*-test used to identify statistically significant differences. Significance levels (*p*-values) indicated in the figure. (**E**) Differential distribution of copy number variations (CNVs) between the two populations.

One hundred individuals were randomly collected from the F_2_ of two populations, respectively. A chromosome number of 200 was detected in all 200 individuals. Whole-genome resequencing of the 200 individuals yielded a total of 126.26 billion 150-bp paired-end reads (18.62 Tb), achieving an average depth of 29.10× per individual (Table S1). After mapping the merged reference genomes of parental goldfish and common carp, we identified a total of 31,718,099 single nucleotide polymorphisms (SNPs) and 11,351,262 insertion and deletion variants (InDels). 30,258,383 SNPs and 10,754,208 InDels were detected in 4nR_2_C_2_-B, while 15,194,956 SNPs and 6,485,632 InDels were found in 4nR_2_C_2_-S (Tables S2-S3). Of these variants, 43.30% of SNPs and 51.88% of InDels were shared by the two populations (Fig. 1C and Fig. S1). The number of SNPs and InDels was higher in 4nR_2_C_2_-B than in 4nR_2_C_2_-S (*t*-test: *p* < 0.001), suggesting higher population diversity for the 4nR_2_C_2_-B population (Fig. 1D). Interestingly, our results revealed a contrasting pattern of genetic diversity for SNPs and InDels across subgenomes. In the 4nR_2_C_2_-B population, genetic diversity was primarily concentrated in subgenome R. Conversely, the 4nR_2_C_2_-S population exhibited higher genetic diversity within subgenome C.

Copy number variation (CNV) is also considered a key genetic variation for allopolyploids. We detected 21,794 CNV events (15,802 in 4nR_2_C_2_-B and 14,192 in 4nR_2_C_2_-S), of which 8,200 (37.6%) were shared among the two populations (Fig. S2 and Tables S3-S4). Similar to the distribution of SNPs, most CNVs (16,513, 75.77%) were detected in subgenome R for the two allopolyploid populations (Fig. S2). Unlike the findings for SNPs and InDels, CNVs were concentrated in subgenome R in both populations (Fig. 1E). Interestingly, although 4nR_2_C_2_-B exhibited higher CNV genetic diversity in subgenome R compared to 4nR_2_C_2_-S, 4nR_2_C_2_-S exhibited higher genetic diversity of CNVs in subgenome C (Fig. 1E). Our above results indicate lower genetic diversity in the 4nR_2_C_2_-S population compared to 4nR_2_C_2_-B.

### Population structure and genetic diversity

To further investigate the genetic diversity between the two populations, we conducted linkage disequilibrium (LD) analysis and observed higher LD values in 4nR_2_C_2_-S compared to 4nR_2_C_2_-B. This observation suggests stronger selective pressures acting on 4nR_2_C_2_-S, which could explain the low genetic diversity observed in this population (Fig. 2A). Principal components analysis (PCA), distance-based neighbor-joining phylogenetic tree, and admixture analysis (K = 2 to 4 clusters) on 67,314,959 SNPs revealed genetic diversity between the 4nR_2_C_2_-B and 4nR_2_C_2_-S populations (Fig. 2B-C and Fig. S3). 4nR_2_C_2_-S exhibited a distinct genetic component compared to 4nR_2_C_2_-B. However, three individuals in 4nR_2_C_2_-B displayed homogeneity with the 4nR_2_C_2_-S population. Meanwhile, analysis of CNVs data reveals a greater overlap of individuals from the two populations compared to the results based on SNPs (Fig. 2B). Consistent with the pattern observed for SNPs, CNV diversity was also higher in the 4nR_2_C_2_-B population compared to the 4nR_2_C_2_-S population (Fig. 2B).

**Fig. 2.**
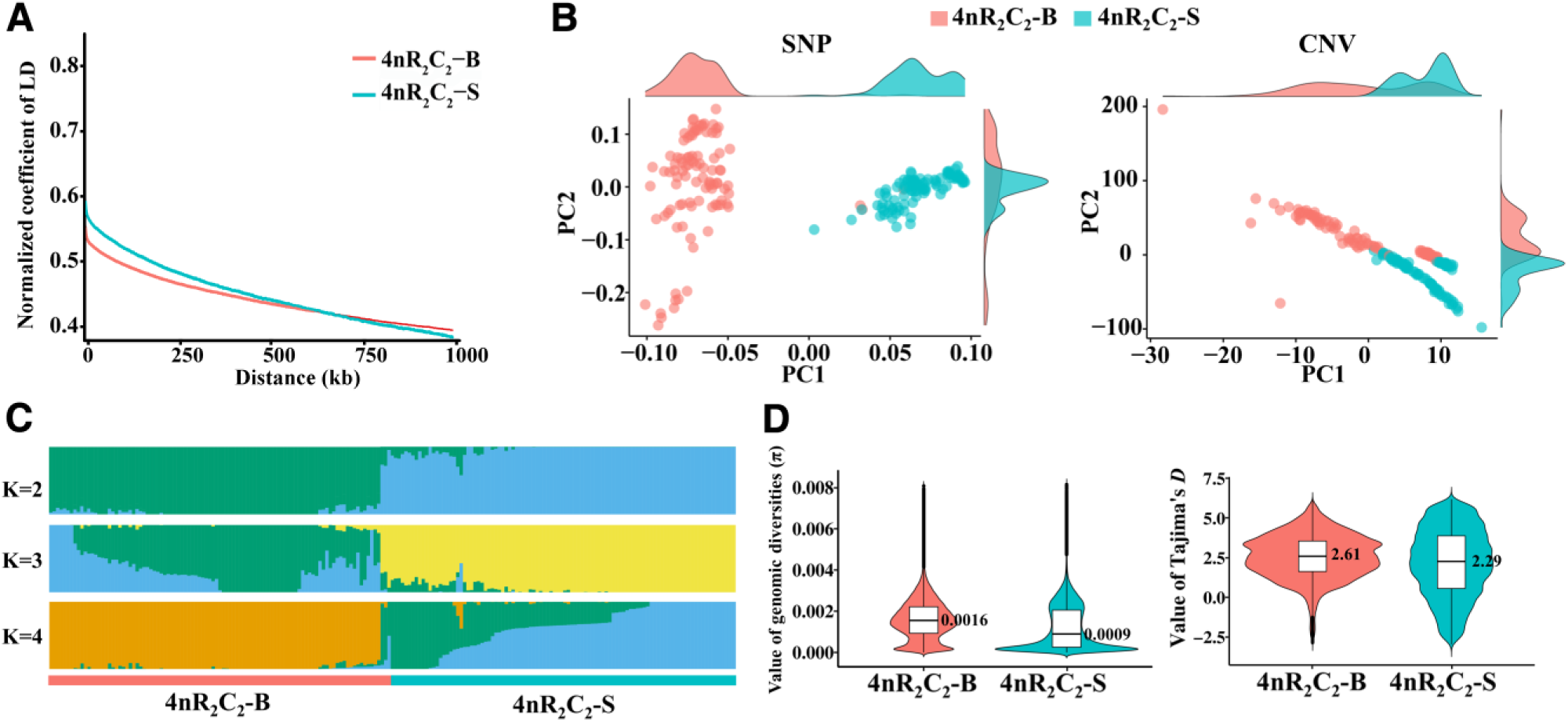
Genetic structure of the two allopolyploid populations. (**A**) Linkage disequilibrium (LD) decays between pairs of SNPs. LD decays relatively slower in 4nR_2_C_2_-S compared to 4nR_2_C_2_-B, where LD decays rapidly and reaches half its maximum value within a shorter physical distance. (**B**) Principal component analysis (PCA) of single nucleotide polymorphisms (SNPs) and copy number variants (CNVs) in two populations. (**C**) Admixture plots generated from whole-genome sequencing (WGS) data. The analysis considers varying numbers of potential ancestral clusters (K), ranging from 2 to 4. Simulations were run with 1000 bootstraps and tenfold cross-validation. Each vertical bar represents a single individual. The bar is divided into colored segments, with the number of segments corresponding to the chosen K value. The length of each colored segment within a bar indicates the estimated proportion of an individual’s genome ancestry that originated from that specific ancestral cluster. (**D**) Genomic diversities (π) and Tajima‘s *D* values in two populations. Significant differences observed between 4nR_2_C_2_-S and 4nR_2_C_2_-B (*t*-test, *p* < 0.001).

Pairwise genome-wide F_ST_ values between 4nR_2_C_2_-B and 4nR_2_C_2_-S indicated a rapid and high level of genomic differentiation between the two populations (average F_ST_ = 0.17). The genomic diversities (π) in 4nR_2_C_2_-B were higher than in 4nR_2_C_2_-S (*t*-test: *p* < 0.001, Fig. 2D). Population genetic test Tajima’s *D* was also higher in 4nR_2_C_2_-B than in 4nR_2_C_2_-S (*t*-test: *p* = 2.30e^-158^, Fig. 2D). Analyses of V_ST_ values based on the 21,794 CNVs indicated a low level of genomic differentiation (average V_ST_ = 0.038) between 4nR_2_C_2_-B and 4nR_2_C_2_-S. 4nR_2_C_2_-S populations exhibit lower levels of genetic diversity, as evidenced by both SNP and CNV data. These findings suggest a decrease in genetic diversity within the 4nR_2_C_2_-S population driven by selection pressure.

### Genomic signatures relate to adaptation

Different environments can impose selection pressures, leading to distinct selective signatures in allopolyploid populations (Monnahan et al., 2019). The above results have demonstrated rapid genetic diversity in SNP levels within two generations for the 4nR_2_C_2_-S population. To gain a deeper understanding of the relationship between these genetic variations and environmental adaptation, we conducted combined analyses of Fst and π values using a 100 kb sliding window and a 10 kb shift across the genome. We obtained 31 putatively selective sweep regions with the top 5% global values (F_ST_ > 0.53, nucleotide diversity (θπ) > 13.31 or < 0.47), which comprised 354 genes, between 4nR_2_C_2_-B and 4nR_2_C_2_-S (Fig. S4). Annotation of these genes revealed functions associated with traits including growth and reproduction (Fig. S5). Functional analysis of the 354 genes revealed significant enrichments for GO categories involving six biological processes such as smooth muscle contraction (GO:0006939), as well as two molecular function categories including growth factor receptor binding (GO:0070851) and fibroblast growth factor receptor binding (GO:0005104) (Table S5).

To investigate positive selection within the 4nR_2_C_2_-S population across two generations, we further analyze genetic diversity in their genomes. We identified 856 (spanned 138.16 Mb) and 397 (spanned 89.33 Mb) selective sweep regions (top 5% values) based on π and Tajima’s *D* analyses, respectively (Tables S6-S7). Functional analysis of the 1192 genes related to selective sweep regions revealed significant enrichments for GO categories involved in 6 biological process categories, including fast-twitch skeletal muscle fiber contraction (GO:0031443) as well as 4 molecular function categories (Table S8). The three pathways, including VEGF signaling pathway (ko04370), were obtained from KEGG pathway analysis (Fig. S6). Our findings suggest a link between adaptation to body size in different water environments and growth-related traits.

### Genome-wide association **s**tudy for growth traits

Growth-regulated genes associated with selective sweep regions and exhibiting differential growth phenotypes were identified in both the two allopolyploid populations and within 4nR_2_C_2_-S (Fig. 1B and Tables S5 and S8). To investigate the relationship between population genetic diversity and growth phenotypes, we conducted a genome-wide association study (GWAS) for growth traits. We found 1,784 SNPs associated with body length and 271 SNPs associated with body height detected in 4nR_2_C_2_-S, while no significant loci was identified in 4nR_2_C_2_-B (Fig. 3A-B, Fig. S7, and Table S9). Selection pressure acting on 4nR_2_C_2_-S could be a key factor in the identification of critical SNPs associated with its growth phenotypes. Interestingly, despite both body length and height being classified as growth phenotypes, the GWAS analysis of the 4nR_2_C_2_-S population revealed only 43 SNPs (2.1%) common to both traits (Fig. 3B). However, a considerably higher proportion (148 genes, or 27.3%) of the genes regulated by these SNPs were shared between the two traits (Fig. 3B and Table S9). Furthermore, the vast majority (over 97% for body length and 70% for body height) of SNPs linked to these traits resided in subgenome R (Table S9).

**Fig. 3.**
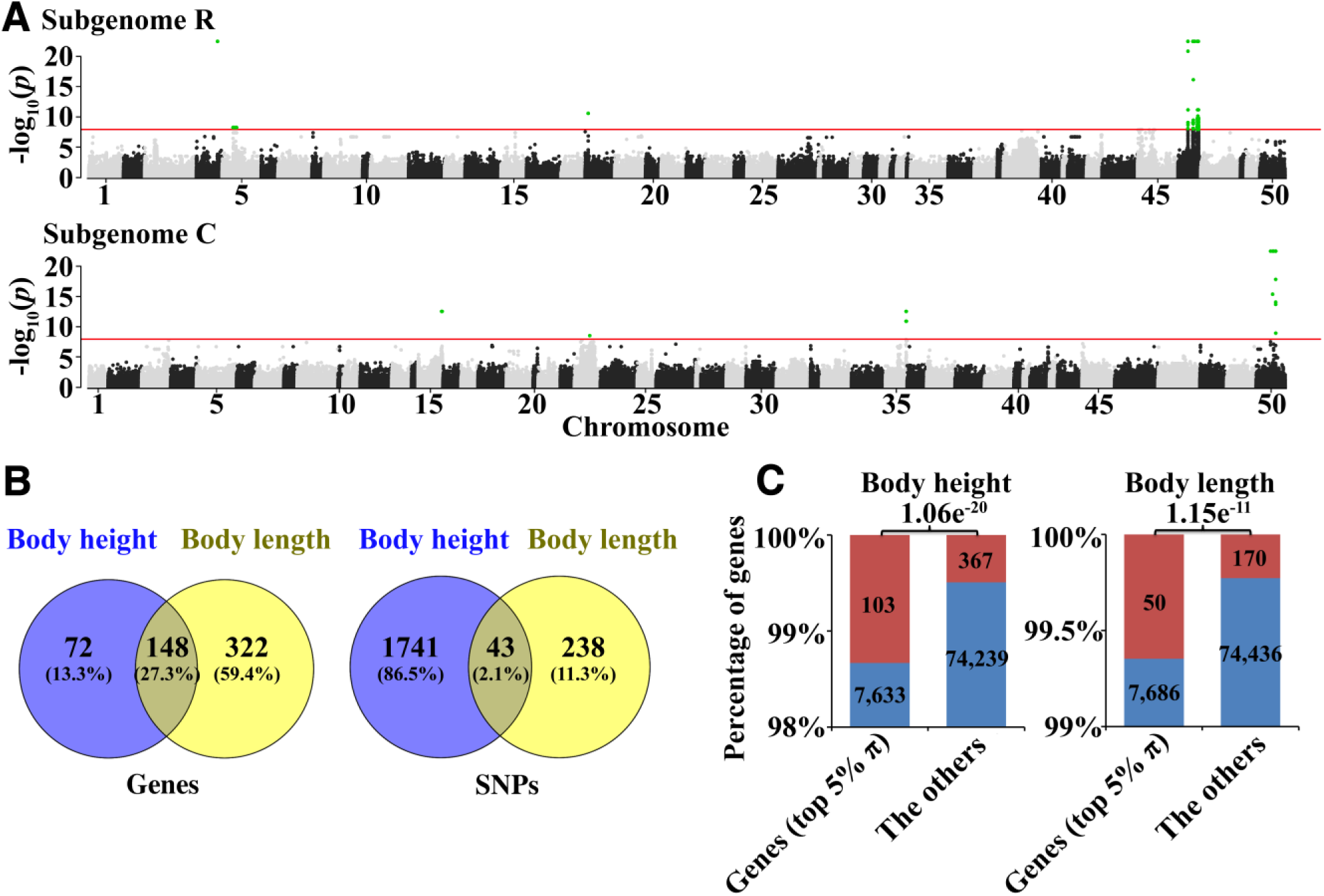
Genome-wide association study (GWAS) of growth traits in 4nR_2_C_2_-S. (**A**) Manhattan plots showing significant SNPs on subgenomes R and C. The x-axis represents the genetic distance along each chromosome, while the y-axis shows the negative base-10 logarithm (log_10_) transformed *p*-values obtained from the GWAS analysis. The red horizontal line represents a suggestive significance threshold of -log10(p) = 8, highlighting SNPs (green dots) exceeding this threshold. (**B**) Distribution of the two growth traits and their association with SNPs and genes. (**C**) Positive correlation between genes under selection (top 5% π values) and growth-regulated genes (Pearson’s chi-square test, *p*-values are signed in figure). Red represents growth-regulated genes, while blue represents the other genes.

Our analysis identified a positive correlation between genes potentially under selection and growth-regulated genes. We compared the 7,770 genes with the highest 5% of π values (indicative of potential selection pressure) to the 470 genes associated with body height and the 220 genes associated with body length identified by GWAS (Pearson correlation: *p* = 1.06e^-20^ and 1.15e^-11^, respectively). This finding suggests that genes under selection may play a role in regulating growth traits (Fig. 3C).

### Gene expression changes in different tissues and organs

To investigate the effects of water body size on various tissues and organs, we collected 66 samples (three biological replicates each from 11 tissues and organs) from two F_2_ allotetraploid populations (Table S10). Gene expression clustering revealed distinct expression patterns across the 11 tissues and organs between the two populations (Fig. 4A and Fig. S8). Analysis of the distribution of gene expression values between 4nR_2_C_2_-B and 4nR_2_C_2_-C populations demonstrated that the intestine and muscle exhibited greater expression differences compared to the other nine tissues and organs (Fig. 4B and Table S11). Similarly, analyses based on Hedges’ g values identified the muscle as having the most significant expression differences (Table S11). Muscle displayed the highest number of differentially expressed genes (DEGs) between populations (1,063 genes, 2.59%) compared to the ovary, liver, heart, kidney, pituitary, and brain. However, the testis, eye, spleen, and intestine exhibited a higher number of DEGs (Fig. 4C and Table S12).

**Fig. 4.**
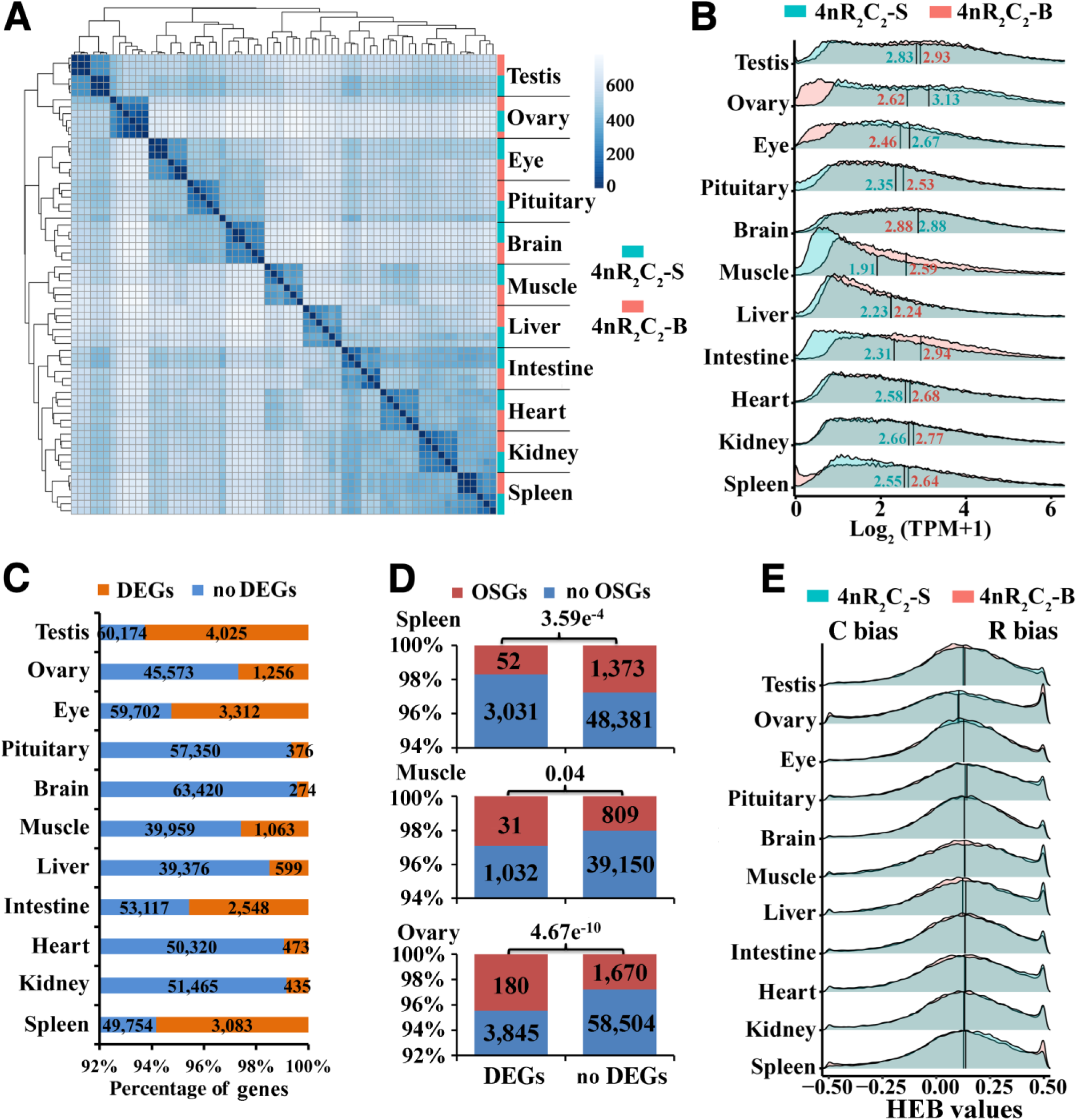
Gene expression profiles across all 11 tissues and organs. (**A**) Organ-specific gene expression. Heatmap showing gene expression levels (color intensity indicates relative expression). (**B**) Population-specific gene expression. Comparison of gene expression between populations (median values in figure). (**C**) Proportion of differentially expressed genes (DEGs) and genes not differentially expressed (no DEGs). (**D**) Overlap of DEGs with organ-specific genes (OSGs) in spleen, muscle, and ovary (*p*-value from Pearson’s chi-square test in figure). Y-axis represents the percentage of genes. *P*-values are signed in figure. (**E**) Homoeologous expression bias (HEB) of the two populations. “R Bias“: higher expression in homoeolog R, “C Bias“: higher expression in homoeolog C.

Deciphering the landscape of organ-specific genes (OSGs) could offer a way to unravel the potential interplay between gene expression, regulatory mechanisms, and physiological processes that govern the diverse functionalities of organs and tissues (Schaum et al., 2020). We detected 1358 OSGs in muscle and found 24 (1.77%) ones shared with DEGs. Furthermore, all the genes shared between the OSGs and DEGs of muscle were the down-regulated genes in 4nR_2_C_2_-B as compared to 4nR_2_C_2_-S (Table S13). In allopolyploids, we classified genes into homologous genes and orphan genes (OGs) to investigate whether the differential expression occurring in different tissues and organs has a specific bias for adaptation to small water bodies (Table S12). Analysis of the allopolyploid’s OGs revealed that 5.58% (2409 out of 43,143) originated from goldfish, and 3.83% (1711 out of 44,625) originated from common carp (Table S12). The proportion of shared and expressed OGs between the two populations was lower in muscle (69.18%) compared to the seven other tissues and organs (brain, intestine, kidney, heart, ovary, testis, and pituitary) (Fig. S9). Interestingly, we observed a positive correlation between differentially expressed genes (DEGs) and OGs in muscle (31 genes) and testis (180 genes), whereas a negative correlation was found in the spleen (52 genes) (Pearson correlation: *p*-value < 0.05) (Fig. 4D and Table S14). The enhanced dynamic changes of OGs in muscle tissue suggest their potential role in adapting muscle development for the allopolyploid’s survival in small water bodies.

Homoeologous expression diversity (HED) is a crucial indicator for evaluating the dynamic changes in gene expression within allopolyploids. We assessed HED for each gene based on the ratio of expression between homoeologs R and C, classifying them into three patterns: homoeolog R bias, homoeolog C bias, and no bias. A higher ratio of homoeolog R bias (ranging from 61.85% to 71.97%) was observed across 11 tissues and organs in both populations (Fig. 4E and Fig. S10). Interestingly, muscle (4nR_2_C_2_-B: 10.71%, 4nR_2_C_2_-S: 10.95%) and liver (4nR_2_C_2_-B: 12.14%, 4nR_2_C_2_-S: 12.28%) exhibited the highest proportion of genes with no bias compared to the other nine tissues and organs. Conversely, the ovary displayed the highest ratio of homoeolog C bias (4nR_2_C_2_-B: 61.85%, 4nR_2_C_2_-S: 62.31%) and the lowest ratio of homoeolog R bias (4nR_2_C_2_-B: 31.19%, 4nR_2_C_2_-S: 30.67%) among all 11 tissues and organs (Fig. S10). Comparing the two populations, the greatest changes in homoeologous expression were observed in the testis (17.28%) and muscle (10.08%) compared to the other nine tissues and organs, which ranged from 2.08% to 8.80% (Table S15). This analysis of dynamic gene expression, considering OSGs, OGs, and HED across different tissues and organs, provides valuable insights into the potential adaptations to a smaller water body size.

### Gene expression atlas in muscle development

The above results demonstrate that both growth phenotypes and growth-related tissues and organs have undergone a series of changes during the adaptation to small water body habitats. To gain deeper insights into the specific growth genes involved in this process, we performed mRNA-seq on muscle samples from 135 individuals (100 from 4nR_2_C_2_-S and 35 from 4nR_2_C_2_-B) (Table S16). Consistent with the findings from the SNP analysis, gene expression clustering revealed that the majority of individuals remained segregated into two distinct groups, despite a small overlap between them (Fig. 5A). Subsequently, we conducted weighted gene correlation network analyses (WGCNA) in each population separately. In 4nR_2_C_2_-B, 299 genes were identified as potential growth-regulated genes, while 426 genes were identified in 4nR_2_C_2_-S (Fig. S11). Notably, only two genes were shared between the two populations, indicating no significant correlation between the genes regulating muscle development in different environments (*p* > 0.05) (Fig. S11). Functional enrichment analysis revealed distinct patterns between the two populations (Fig. S12). Genes identified in 4nR_2_C_2_-B were primarily enriched for skeletal system development (GO: 0001501), potentially reflecting a prioritization of structural growth in this population. Conversely, genes from 4nR_2_C_2_-S exhibited enrichment in cellular energy metabolism pathways (GO: 0006091 and GO: 0019646). This enrichment suggests an adaptation for efficient energy utilization, potentially crucial for survival in the small water body environment.

**Fig. 5.**
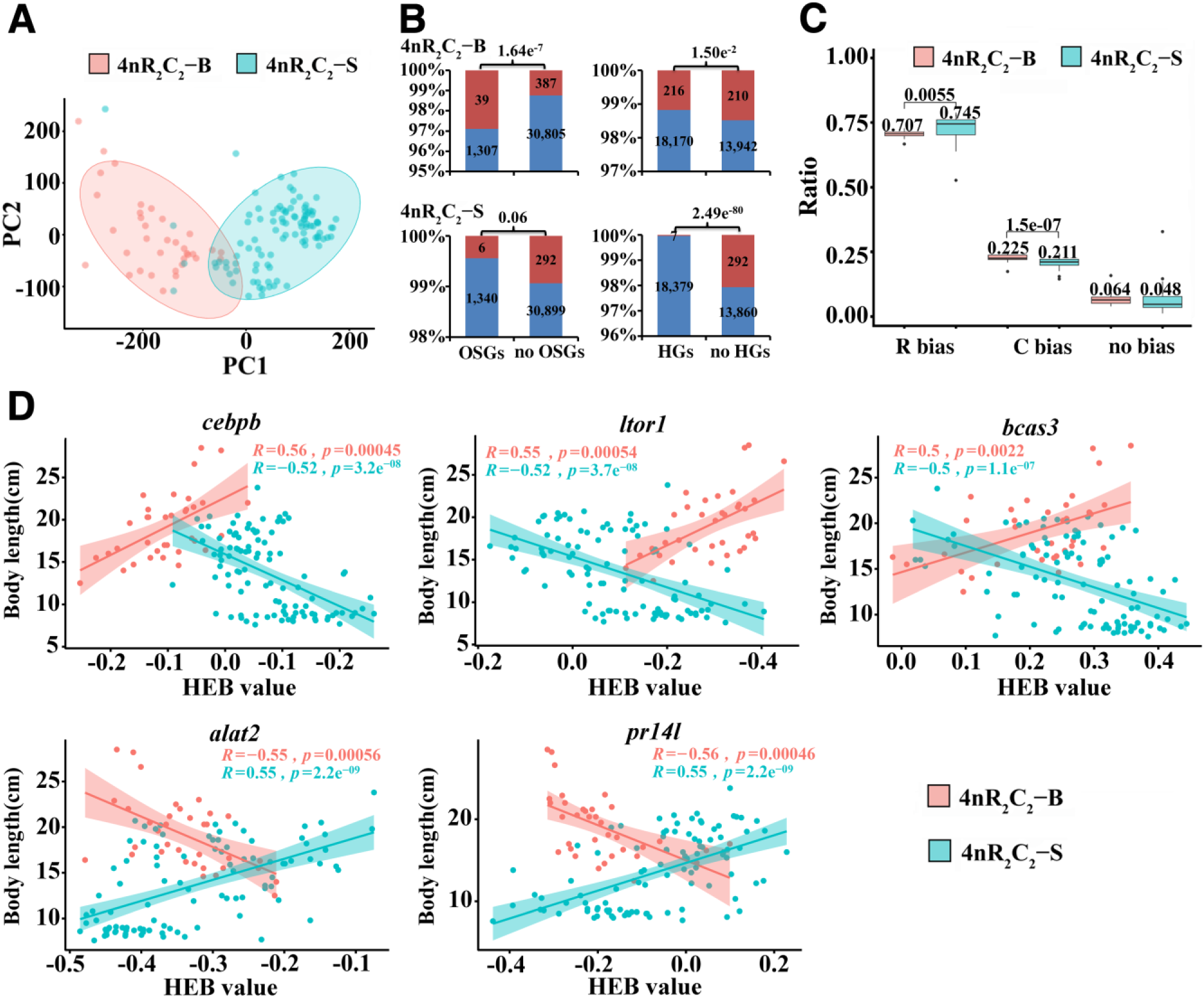
Gene expression profiles in muscle tissue (135 individuals). (**A**) Principal component analysis (PCA) plot of gene expression variation between the two populations based on muscle tissue data. (**B**) Pearson’s correlation coefficients between two gene sets (organ-specific genes (OSGs) or homoeologous genes (HGs)) and growth-related genes in muscle tissue. “OSGs” represents genes specifically expressed in muscle, while “no OSGs” represents genes not specifically expressed in muscle. Similarly, “HGs” represent genes with homoeologs, and “no HGs” represents genes without homoeologs. Red represents growth-regulated genes, while blue represents the other genes. Y-axis represents the percentage of genes. *P*-values are signed in figure. (**C**) Proportion of genes exhibiting the three homoeologous expression bias patterns (“R Bias“—higher expression in homoeolog R, “C Bias“—higher expression in homoeolog C) in the muscle tissue. (**D**) Pearson’s correlation coefficients between the values of homoeologous expression bias (HEB) and body length (cm) in five genes (*cebpb, ltor1*, *bcas3*, *alat2*, and *pr14l*).

Our overall analysis revealed contrasting regulatory mechanisms for growth regulation following polyploidization in 4nR_2_C_2_-S and 4nR_2_C_2_-B populations (Fig. 5B, Fig. S13). In 4nR_2_C_2_-S, a positive correlation between growth-associated genes and muscle-specific genes suggests a dominant role for muscle growth in regulating body size within this population. Conversely, the absence of a significant correlation in 4nR_2_C_2_-B implies alternative regulatory pathways, potentially less reliant on specific organ expression. Furthermore, the negative correlation between growth-associated genes and homoeologous genes (HGs) in both populations (Fig. 5B) suggests subfunctionalization or neo-functionalization of these genes, leading to diverse roles in growth regulation. Notably, all six OGs in the 4nR_2_C_2_-B population showed positive correlations with growth, while all three OGs in the 4nR_2_C_2_-S population displayed negative correlations (Fig. S13). These contrasting patterns suggest population-specific adaptations in growth regulation strategies, potentially linked to the smaller water body size experienced by 4nR_2_C_2_-S.

To investigate the effects of homoeologous gene expression bias on growth, we performed correlation analyses between homoeologous expression bias values and body length. Consistent with the findings in multiple tissues and organs, a bias towards the R homoeolog was detected in most (52.62%-78.89%) of the 6,940 expressed homoeologous genes across all individuals (Fig. 5C). Notably, 4nR_2_C_2_-B individuals exhibited a significantly higher R homoeolog bias and lower C homoeolog bias compared to 4nR_2_C_2_-S individuals (*t*-test: *p* < 0.01). 4nR_2_C_2_-S exhibited a significantly higher proportion of genes (359), displaying a correlation between homoeologous expression bias and body length compared to 4nR_2_C_2_-B (95 genes) (Pearson correlation: *p*-value < 0.05, r > 0.5 or r < −0.5) (Fig. S14). This finding implies that the dynamic interplay of homoeologous expression bias across multiple genes may facilitate body size adaptation in allopolyploids when faced with environmental shifts. Interestingly, five genes (*cebpb, ltor1*, *bcas3*, *alat2*, and *pr14l*) exhibited a reversal of the correlation coefficient pattern between the two populations. This observation suggests that homoeologous expression bias may exert contrasting effects on individual growth under different aquatic environmental conditions (Fig. 5D and Table S17). Overall, our results highlight the distinct gene expression patterns and regulatory mechanisms underlying growth regulation in the two populations. The observed differences potentially reflect adaptations to the contrasting environmental challenges faced by each population, particularly the smaller water bodies inhabited by 4nR_2_C_2_-S.

### Epigenetic basis of water body size diversification

To investigate the epigenetic basis underlying the diversification in body size observed between the 4nR_2_C_2_-B and 4nR_2_C_2_-S populations, we explored changes in chromatin accessibility, a key epigenetic mark associated with gene regulation. We performed ATAC-seq to profile chromatin accessibility in muscle samples from both populations, mirroring the tissues and organs used for mRNA-seq analysis. The high sequencing depth and preferential mapping to transcription start and end sites (TSS/TES) indicate robust data quality (Tables S18-S20 and Fig. S15). Comparative analysis between 4nR_2_C_2_-B and 4nR_2_C_2_-S revealed a relatively small subset (1.74%, 11,201 regions) of differentially accessible chromatin regions (DARs) across the entire genome (Table S21). This observation suggests that most chromatin accessibility patterns are conserved between the populations. Furthermore, similar proportions of DARs were observed within each subgenome (R and C), further highlighting the overall similarity in the chromatin landscape (Fig. 6A). DARs were more frequently found in distal intergenic regions compared to proximal promoter regions, suggesting their potential role in regulating non-coding elements (Table S21).

**Fig. 6.**
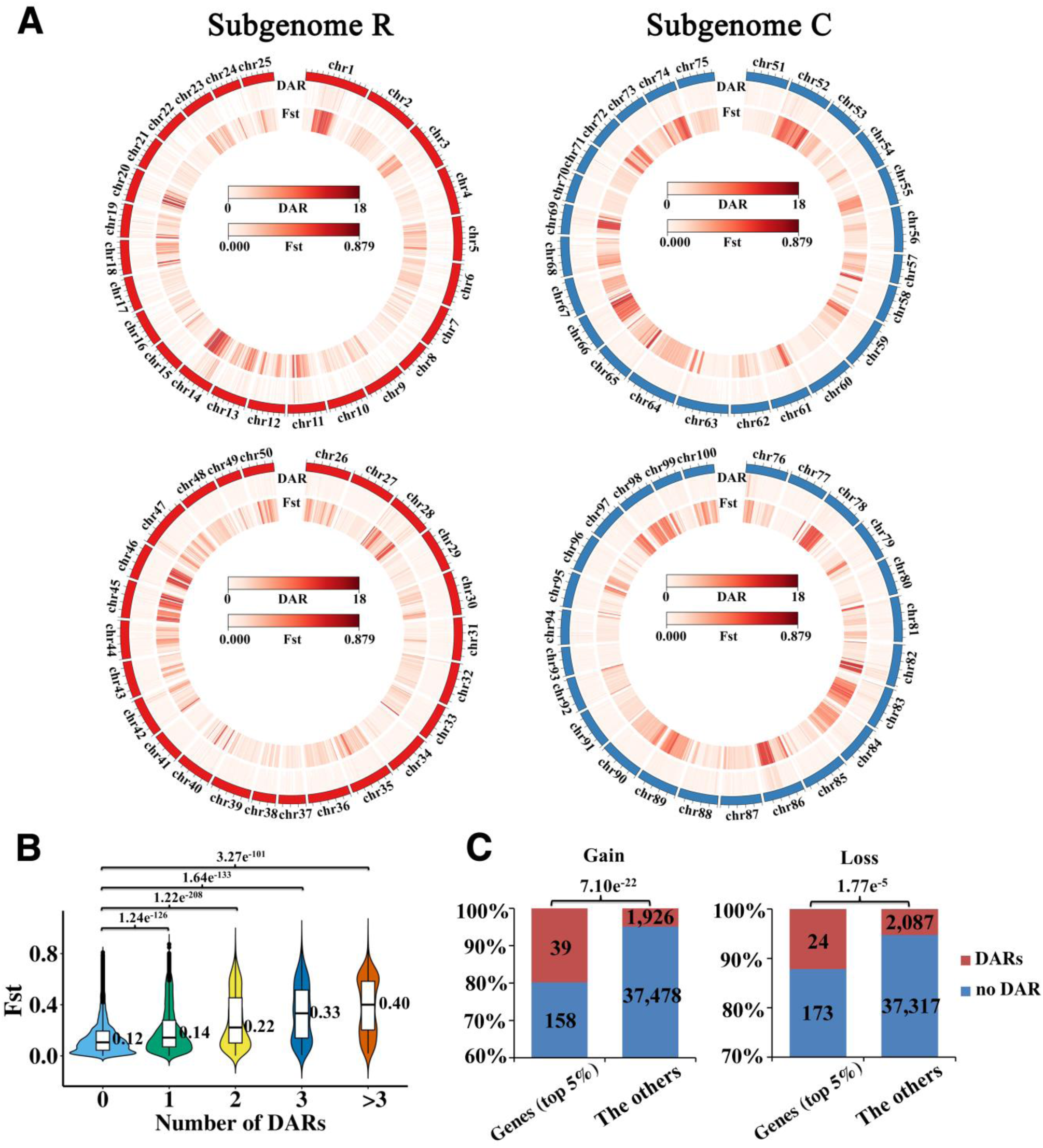
Relationship between accessible chromatin and genetic diversity. (**A**) A circle plot exhibiting the Fst value and DARs number across 100 chromosomes. (**B**) Statistically significant differences (*t*-test: *p* < 0.001) between differentially accessible regions (DARs) with zero associated genes and those with one or more associated genes (>= 1) in the two populations. (**C**) Positive correlations between selection genes (top 5% based on Fst and π values) and DAR-associated genes, identified using Pearson correlation analyses. “Gain” represents increased chromatin accessibility in 4nR_2_C_2_-S than in 4nR_2_C_2_-B. Conversely, “Loss” represents decreased chromatin accessibility in 4nR_2_C_2_-S. “DARs” represents DAR-associated genes, while “no DARs” represents the other genes. Y-axis represents the percentage of genes. *P*-values are signed in figure.

We identified 8,023 genes associated with DARs, potentially under the regulatory influence of these accessibility changes (Fig. S16). A positive correlation was observed between the degree of genetic diversity (Fst) and the number of DARs across the genome (Fig. 6B). This suggests a potential link between regions experiencing selection pressure (reflected by Fst) and chromatin accessibility changes. Further analysis of the top 5% most differentiated genes (based on Fst and π values) revealed a positive correlation with DAR-associated genes located on the R subgenome (Fig. 6C). This finding suggests that subgenome R might be more susceptible to chromatin accessibility changes associated with genetic diversity.

Joint analysis of ATAC-seq and RNA-seq data revealed a mixed relationship between chromatin accessibility and gene expression. Some genes (58) and their associated DARs (89) displayed a positive correlation, suggesting that increased accessibility might enhance gene expression. Conversely, a negative correlation was observed in another subset of genes (47) and DARs (59) (Fig. S18). These contrasting findings highlight the complex interplay between chromatin accessibility and gene regulation. Further investigation into the specific functions of these genes could provide valuable insights into the regulatory pathways underlying water body size diversification.

## Discussion

Allopolyploidization, a powerful evolutionary force, has significantly shaped the genetic and phenotypic diversity of fish species such as goldfish, common carp, and gibel carp (*Carassius gibelio*) influencing traits like caudal fin number, body height-to-length ratio, and coloration. Moreover, these polyploid individuals exhibit remarkable resilience to phenotypic mutations, rarely succumbing to even substantial morphological alterations. This observation underscores the significance of allopolyploidization as a driver of rapid phenotypic diversification and adaptive evolution. This study explores the interplay between allopolyploidization and water body size in sculpting body size evolution. We utilize a nascent allopolyploid fish population (4nR_2_C_2_, derived from goldfish and common carp) as a model system to illuminate the underlying mechanisms of adaptation to different water environments.

Geological and monsoonal natural environmental changes, as well as human activities, often lead to rapid alterations in the structure, morphology, and water quality of rivers and lakes worldwide (Dodds, 2006). These alterations can lead to short-term shifts in the diversity and abundance of aquatic organisms, encompassing both plankton and fish species (Mao et al., 2021). However, over time, the dissemination of adaptive mutations within populations empowers aquatic organisms equipped to adapt to the transformed environment to proliferate, ultimately leading to a metamorphosis of community structure and the emergence of novel species (Lloyd, Makukhov, & Pespeni, 2016). Our study reveals evidence of selective sweep and high genetic homogeneity in an allopolyploid fish population inhabiting a small water body with a harsh environment. This population exhibits significant genetic differentiation from another population residing in a larger water body, and this differentiation occurred within just two generations. This rapid evolutionary adaptation may be linked to the unique genetic regulatory mechanisms of allopolyploidy.

This diversity is manifested in distinct patterns of SNPs and CNVs across the subgenomes. Notably, the high enrichment of genetic variation within subgenome R of 4nR_2_C_2_-B suggests a potential link to stronger selection pressures or a higher propensity for rearrangements within this subgenome in response to environmental cues. This finding aligns with the phenomenon of maternal dominance observed in allopolyploid cyprinid fishes (M.-R.-X. Xu et al., 2023). Population genetic analyses further substantiate this differentiation. The lower genetic diversity and higher population structure in 4nR_2_C_2_-S compared to 4nR_2_C_2_-B suggest rapid genetic changes driven by unique selective forces within the smaller water body. Notably, selective sweep regions associated with growth-related traits highlight ongoing adaptation to water body size. Functional enrichment analysis of genes within these regions reinforces the centrality of growth and metabolism pathways in environmental adaptation.

We delve deeper into the regulatory mechanisms governing growth in response to water body size. Genome-wide association studies (GWAS) pinpoint candidate loci associated with body length and height, revealing population-specific patterns. Additionally, the differential expression of growth-related genes across different tissues and organs emphasizes the role of their specific adaptations. Interestingly, homoeologous gene expression bias emerges as a potential mechanism influencing growth, with population-specific patterns suggestive of adaptive responses to environmental cues. Epigenetic analyses reveal changes in chromatin accessibility linked to the genetic diversity between populations. This interplay between genetic and epigenetic factors likely regulates gene expression in response to environmental cues.

This study also sheds light on the evolutionary trajectories of goldfish and common carp. Previous research revealed intriguing subgenome divergence: goldfish subgenome P undergoes pseudogenization, suggesting a distinct evolutionary strategy compared to common carp subgenome A (D. Chen et al., 2020; Z. Chen et al., 2019; Kon et al., 2020; Luo et al., 2020). This subgenome-specific flexibility likely facilitates adaptation to diverse environments and contributes to phenotypic diversification (Z. Li et al., 2023; Peng et al., 2022; M.-R.-X. Xu et al., 2023). Additionally, the striking similarity in expression bias towards matrilineal homoeologs hints at a potential link to water body size adaptation. Goldfish, which have a higher maternal bias and smaller body size, are more likely to thrive in smaller environments, while common carp, which have a stronger paternal bias and larger body size, are more likely to inhabit larger water bodies (Luo et al., 2020; P. Xu et al., 2014). This pattern suggests a complex interplay between subgenome evolution, gene expression, and adaptation to varying water body sizes in these polyploid fish.

Our study uncovers the critical role of allopolyploidization in fish body size evolution and adaptation to water body size. We demonstrate the rapid adaptive potential of allopolyploid fish, accompanied by rapid genomic divergence and epigenetic modifications. These findings provide significant insights into polyploid speciation and differentiation. Nevertheless, future research focusing on the impact of multigenerational genetic and epigenetic variation on body size evolution, functional validation of candidate genes and epigenetic modifications, the ecological significance of body size variation in natural allopolyploid fish populations, and comparative studies across diverse allopolyploid systems and environmental gradients will be crucial to further elucidating the role of allopolyploidization in fish body size evolution and adaptation, as well as the mechanisms underlying the adaptation and differentiation of polyploid species in natural environments.

## Materials and Methods

### Study design

To investigate how the nascent allotetraploid fish, 4nR_2_C_2_ (derived from goldfish [*Carassius auratus*] and common carp [*Cyprinus carpio*] hybridization), adapts to different water body sizes, we established two experimental pools with contrasting sizes in Changsha, Hunan Province, China (28°11′ N, 112°58′ E). Both pools (small: 8.8 m × 4.7 m × 1.4 m; large: 50.0 m × 65.0 m × 2.5 m; Fig. S) maintained similar environmental conditions, including illumination, water temperature, adequate food supply, and appropriate oxygen content. One thousand 4nR_2_C_2_ fry were randomly stocked into each of four ponds (two small, two large) in May 2020. These fish become sexually mature at one year (Shaojun Liu et al., 2001). Following the breeding season, adult fish were removed from both pools, and the resulting fry were raised until the next breeding season in June 2021. In April 2022, a random sample of 50 healthy 12-month-old juvenile fish was collected from each pool, totaling 100 fish per pool size.

### Sampling and ploidy determination

Before tissue collection, the ploidy level of each individual was identified using flow cytometry. The DNA content of erythrocytes of *C. auratus* (goldfish, 2nRR), *C. carpio* (common carp, 2nCC), and 4nR_2_C_2_ was measured using flow cytometry (Cell Counter Analyzer, Partec, Germany). About 0.5–1 ml of blood from each fish was collected from the caudal vein using a syringe containing 100–150 units of sodium heparin. The blood samples were treated following the method described in (Xiao et al., 2014). The DNA contents of 2nRR and 2nCC were used as controls. These fish were deeply anesthetized with 300 mg/L Tricaine Methanesulfonate (Sigma-Aldrich, St. Louis, MO, USA) for 10 min (25°C) in a separate tank. After confirming death, they were collected for dissection. Some growth traits, including body length, body height, height of back muscle, and body weight, were detected for each individual.

### DNA isolation and whole genome re-sequencing

To investigate the relationship between genomic variations and body size involving different growth environments in the allopolyploid fish, high-quality genomic DNA of the muscle from 4nR_2_C_2_ was isolated using the DNeasy Blood & Tissue Kits (Qiagen) method. The quality of DNA was checked by a NanoDrop^®^ ND-1000 spectrophotometer with a 260/280 ratio and a 260/230 ratio. DNA integrity and purity were checked using gel electrophoresis. Whole genome sequencing was performed with DNA nanoball (DNBSEQ-T7) technology. The three stages were performed in the whole genome sequencing: 1) the construction of single-stranded circular libraries with a paired-end library (150 bp × 2); 2) the generation and loading of DNBs onto patterned nanoarrays; and 3) combinatorial probe anchor synthesis sequencing (Patterson et al., 2019). The raw data was subjected quality checking and adapter removal using Fastp (v. 0.21.0) (S. Chen, Zhou, Chen, & Gu, 2018). The high-quality reads were used in the next analysis.

### SNP calling from genomic data

For whole-genome sequencing data, raw paired-end reads were filtered by Fastp (v. 0.21.0) and then aligned to the merged files of goldfish (Luo et al., 2020) and common carp (P. Xu et al., 2019) genomes using BWA (v. 0.7.17-r1188) (Kim, Paggi, Park, Bennett, & Salzberg, 2019). Aligned reads were converted to BAM format using Samtools (v. 1.16), and duplicate reads were marked with the MarkDuplicates method of Picard (v. 2.27) (http://broadinstitute.github.io/picard). Next, HaplotypeCaller, CombineGVCFs, GenotypeGVCFs, SelectVariants, and VariantFiltration in the Genome Analysis Toolkit (GATK, v. 4.0.4.0) pipeline were used to call, filter, and select single nucleotide polymorphisms (SNPs) and small insertions and deletions (InDels). The filter parameters were as follows: “QUAL < 30.0 || QD < 2.0 || MQ < 40.0 || FS > 60.0 || MQRankSum < −12.5 || ReadPosRankSum < −8.0 -clusterSize 2 -clusterWindowSize 5”.

### Population structure analysis

To construct a phylogenetic tree, we used SNPs with a minor allele frequency (MAF) below 0.05 and integrity (INT) greater than 0.8. The neighbor-joining method and the p-distance model were implemented in MEGA X for this purpose. The final consensus tree was validated using bootstrapping with 1,000 replicates. Population structure analysis was performed using ADMIXTURE (v. 1.22) based on the filtered SNPs. We evaluated different numbers of subpopulations (K) ranging from 1 to 10. Each K value was cross-validated to identify the optimal number of clusters that minimizes the cross-validation error rate. EIGENSOFT (v. 6.0) was employed for principal component analysis (PCA) based on the SNP data. Linkage disequilibrium (LD) between pairs of SNPs within a 1,000 kb window on the same chromosome was calculated using PopLDdecay (v. 3.41). The normalized coefficient of LD (*R*²) was used to quantify the correlation between loci. LD decay maps were constructed for each genome and the entire genome by plotting *R*² values against genetic distance (bp). The LD decay distance is defined as the distance at which *R*² is reduced to half of its maximum value.

### Detection of genomic signatures of selection

To assess population differentiation between and within populations, we employed four indices: Fst (fixation index) for genetic divergence, Tajima’s *D*, nucleotide diversity (π), and CNV analysis using Vst (variance stabilizing transformation, a population differentiation metric similar to Fst). A window size of 100 kb with a 10 kb step size was used to investigate Fst, Tajima’s *D*, and π. Tajima’s *D* compares nucleotide diversity (θπ) to identify deviations from the expected neutral site-frequency spectrum. These indices were calculated using the vcftools package based on pre-defined bins and steps for consensus SNPs. Genomic windows with the top 5% highest Fst values were considered significant and analyzed for gene function. Gene Ontology (GO) term enrichment analysis was performed, with a significance threshold of *p*-value ≤ 0.05.

### Copy number detection

To identify whether copy number (CN) was also divergent between the two populations, we first identified windows showing significant variation in normalized read depth. Here, we refer to CNV of 0 as an “absence” rather than a deletion because the absence of a locus in an analyzed genome could be the result of a gain in the reference genome and not solely a deletion in the analyzed genomes. To screen genomes for CNVs, we used the *cn.MOPS* R package with the *cn.mops* algorithm (the default bin size was 1 kbp), which is specifically designed for diploid organisms. We calculated Vst based on a range of 0 (no difference in CN allele frequencies between the two populations) to 1 (complete differentiation of CN allele frequencies between the two populations). The formula for Vst is as follows:

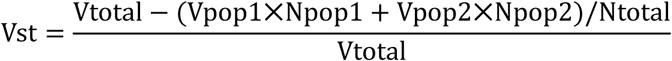

“Vpopx” is the CN variance for each respective population, “Vtotal” is the total variance, “Npopx” is the sample size for each respective population, and “Ntotal” is the total sample size. We considered Vst values in the upper 95^th^ percentile as significantly different between populations.

### Genome-wide association study

We conducted a genome-wide association study (GWAS) to identify genes associated with three growth traits (body weight, length, and height) in 4nR_2_C_2_-B and 4nR_2_C_2_-S populations. Firstly, we removed SNPs with a minor allele frequency (MAF) greater than 0.05 and integrity (INT) greater than 0.8. This filtering step helps to focus on high-quality SNPs that are more likely to be associated with the traits. Then, we addressed potential population stratification by performing principal component (PC) and kinship matrix analyses. The first three principal components (PCs) were used to construct a PC matrix, which was then included in the GWAS model to account for population structure. Furthermore, we performed GWAS using FaST-LMM (factored spectrally transformed linear mixed models) with default settings. We also compared this approach with EMMAX and LMM algorithms to assess the consistency of results across different methods. To identify statistically significant associations, we calculated a threshold using the formula “-log10(1/effective number of independent SNPs)”. This formula accounts for multiple tests and adjusts for the number of SNPs analyzed. Based on this calculation, a significance threshold of -log(P) = 8 was set. Lastly, we visualized the results using Manhattan plots to identify candidate loci exceeding the significance threshold. Additionally, quantile-quantile (QQ) plots were used to assess potential biases in the data and ensure the reliability of the association signals. To understand the biological functions of the genes associated with the identified loci, we performed GO term enrichment analysis. This analysis identified groups of genes with enriched GO terms (*p*-value ≤ 0.001), providing insights into the biological pathways potentially underlying the variation in growth traits.

### RNA isolation and transcriptome sequencing

Total RNA of the 11 tissues and organs, including intestine, spleen, liver, heart, kidney, pituitary, brain, eye, ovary, testis (three biological replicates in each allopolyploid population), and muscle (100 individuals from 4nR_2_C_2_-S population and 35 individuals from 4nR_2_C_2_-B population) was isolated and purified according to a TRIzol extraction method (Rio, Ares, Hannon, & Nilsen, 2010). RNA concentration was measured using NanoDrop technology. Total RNA samples were treated with DNase I (Invitrogen) to remove any contaminating genomic DNA. The purified RNA was quantified using a 2100 Bioanalyzer system (Agilent, Santa Clara, CA, USA). 1 µg of isolated mRNA was fragmented with fragmentation buffer. Paired-end (2 × 150 bp) sequencing was performed using DNA nanoball (DNBSEQ-T7) technology according to the standard method (Patterson et al., 2019). Then, low-quality bases and adapters were trimmed out using Fastp (v. 0.21.0). The high-quality reads were used in the next analysis.

### Expression calculation and differential expression analysis

All mRNA-seq reads were mapped to the combined nuclear and mitochondrial genomes of *C. auratus* (Luo et al., 2020) and *C. carpio* (P. Xu et al., 2019) using BWA (v. 0.7.17-r1188) (Kim et al., 2019) with default parameters. Then, the mapped files were handled with Samtools (v. 1.10) (H. Li et al., 2009), while the unique mapped reads were obtained using htseq-count (Srinivasan, Virdee, & McArthur, 2020). The expression value was normalized based on the ratio of the number of mapped reads for each gene to the total number of mapped reads for the entire genome. The transcripts per million (TPM) values were calculated based on the normalized data. These reads were used to calculate the expression values of alleles R (origin from goldfish) and C (origin from common carp) in allopolyploid (L. Ren et al., 2017). Differential expression analysis was performed with the thresholds (fold change > 3, *p*-value < 0.001, and padj < 0.001) using Deseq2 of R package. Expressed genes were determined based on the thresholds, with read counts in each sample > 9 and TPM > 0.5. Hedges’ g value was used to assess the effect size in different tissues and organs between the two populations (Strauss, Cavanagh, Oliver, & Pettman, 2014).

### Determination of homoeologous, organ-specific, and orphan genes

In an allopolyploid individual, genes can also be classified as homoeologous genes (with orthologs present in both parental species) or orphan genes (present only in one parental species). Genes can be classified into organ-specific genes based on their highest expression among all tissues and organs. Homoeologous gene of the subgenomes R (originating from 2nRR) and C (originating from 2nCC) in the allopolyploid were obtained using the all-against-all reciprocal BLASTP (v 2.8.1) with an e-value cutoff of 1e ^−6^ based on protein sequences (sequence alignment > 70%). Then, transcripts that were shorter than 300 bp were discarded from homoeologous genes. Orphan genes (OGs) of the two species are detected based on the thresholds of BLASTx with an e-value of 1e ^−5^ and tBLASTx with an e-value of 1e ^−5^. The sequences with no BLAST result in the public database were considered potential OGs. Then, the expressed OGs (TPM > 10) were considered OGs in the corresponding tissues and organs. Organ-specific genes (OSGs) were identified using the following criteria: Gene expression in the target tissue or organ differs significantly from that in the other ten tissues and organs. GO analysis was performed with a significance threshold (false discovery rate of Benjamini–Hochberg method < 0.05).

To analyze the diversity in expression levels between homoeologs R and C, genes with mapped read counts exceeding 9 for both homoeologs were selected. We then classified these genes into three expression patterns based on significant (*p*-value < 0.05) deviations from equal expression (50/50) between the homoeologs, determined using a binomial test. HEB values were calculated using the formula: R/(R + C) - 0.5. R represents the reads number of homoeolog R, while C represents the reads number of homoeolog C. These patterns are: Homoeolog R Bias: Genes with significantly higher expression in homoeolog R compared to C, as indicated by a positive “Bias effect” value. Homoeolog C Bias: Genes with significantly higher expression in homoeolog C compared to R, as indicated by a negative “Bias effect” value. No Bias: Genes showing no statistically significant difference in expression levels between homoeologs R and C (“Bias effect” not significant).

### Weighted gene correlation network analysis (WGCNA) and functional annotation

To investigate the relationship between expression patterns and growth traits, we conducted coexpression analysis across 4nR_2_C_2_-S and 4nR_2_C_2_-B populations using WGCNA (v. 1.72). An unsupervised network was constructed based on gene expression originating from different parents using default parameters as follows (Langfelder & Horvath, 2008): First, a matrix of Pearson correlations between genes was generated based on expression values. Then, an adjacency matrix representing the connection strength among genes was established by raising the correlation matrix to a soft threshold power. Next, the adjacency matrix was used to calculate a topological overlap matrix (TOM). Genes with similar coexpression patterns were clustered using hierarchical clustering of dissimilarity. Pearson correlations between the expression level of that gene and module were performed using eigengene based connectivity, while Pearson correlations were further calculated to measure the strength and direction of association between modules and the growth traits (body weight, body length, and body height) (*p*-value < 0.01). Coexpressed modules (5 modules in 4nR_2_C_2_-B and 6 modules in 4nR_2_C_2_-S) were determined and used to in next functional analyses. Then, the potential growth-regulated genes were obtained from these coexpressed modules based on a *p*-value < 0.5 in three growth traits. For the genes in each module, functional enrichments were conducted and annotated with GO and KEGG databases.

### ATAC-seq data generation and processing

The six muscle samples (three biological replicates in 4nR_2_C_2_-S and 4nR_2_C_2_-B) were the same as the above mRNA-seq analyses that were performed with ATAC-seq. Tissues were initially washed with a 0.09% NaCl solution, followed by cryogenic grinding into powders. The powders were then treated with lysis buffer and incubated for 10 minutes at 4 ℃ on a rotation mixer. The resulting cell suspension underwent filtration through a 40 μm cell strainer and was washed three times with cold PBS buffer. After verifying nuclei purity and integrity through microscopic examination, approximately 50,000 nuclei were allocated for tagmentation following established protocols (Corces, Trevino, Hamilton, Greenside, & Sinnott-Armstrong, 2017). Tn5-transposed DNA fragments were purified using AMPure DNA magnetic beads. A subset of the DNA underwent qPCR to determine the optimal number of PCR cycles (average 11). The amplified libraries were analyzed on an Agilent Tapestation 2200 (Agilent Technologies) with a D5000 DNA ScreenTape to assess quality by visualizing nucleosomal laddering. All ATAC experiments were conducted with biological replicates performed in duplicate. The final library was sequenced on the Illumina HiSeq X Ten platform (San Diego, CA, United States) in 150 PE mode.

After obtaining raw data from ATAC-seq, sequencing quality control was performed using cutadapt (http://cutadapt.readthedocs.io/en/stable) (v. 1.9.1). Clean reads were aligned to the merged genomes of goldfish (Luo et al., 2020) and common carp (P. Xu et al., 2019) using Bowtie2 (v. 2.2.6) (Langmead & Salzberg, 2012). Reads mapping to the mitochondrial genome were eliminated using removeChrom (https://github.com/jsh58/harvard). Duplicates were removed using Picard (v. 1.126) (http://broadinstitute.github.io/picard/). Two important measures were used to judge the quality of the ATAC-seq data: the insert size distribution of sequenced fragments and the enrichment score around Transcription Start Sites (TSS). Following library normalization, the signal value at the center of the insert size distribution was used as the TSS enrichment metric.

To find peaks with a high level of confidence in biological replicates, we used MACS2 (--nomodel --extsize 200 --shift −100) for peak calling on biological replicates (Zhang et al., 2008). Peaks identified in three biological replicates were considered to have high-to-medium confidence. Then, we performed an irreproducible discovery rate analysis with a stringent threshold of 0.05 on peaks among three biological replicates. We merged the peaks from all samples to create a consensus peak set. We identified statistically significant differentially accessible regions (DARs) utilizing the R package DESeq2, employing a false discovery rate (FDR) threshold of less than 0.05, and considering peaks with a log2 fold change (FC) exceeding 1. Motif analysis of DARs was performed using the MEME suite with default settings (*p*-value < 0.01) (Heinz et al., 2010). Gene ontology enrichment analysis was conducted using the R package ‘ClusterProfiler’. All sequencing tracks were visualized with the Integrated Genomic Viewer (IGV, v. 2.3.61).

## Data availability

Genomic sequencing data for 200 individuals and mRNA-seq data for 135 individuals have been deposited in the National Genomics Data Center (NGDC) under the BioProject (https://ngdc.cncb.ac.cn/bioproject/browse/PRJCA024756) with accession number CRA015751 and CRA015944.

## Authors’ Contributions

L.R. wrote the manuscript. S.J.L. and L.R. modified the manuscript and designed the study. H.D., L.R., Y.Y.Z., S.F.H., and M.X.L. carried out bioinformatics analyses. Y.K.T., X.G., J.L.C., X.Y.Z., H.Z., M.D.L., L.L., H.L.Z., K.K.L., and S.W. performed aquaculture experiments and extracted raw material. All authors read and approved the final manuscript.

## Funding

This research was supported by National Natural Science Foundation of China (32293252, 32341057, and U19A2040), National Key Research and Development Plan Program (2023YFD2401602), Special Funds for Construction of Innovative Provinces in Hunan Province (2021NK1010), Earmarked Fund for China Agriculture Research System (CARS-45), the 111 Project (D20007), and The world-class cultivation discipline (Biology) of Hunan province.

## Ethics approval and consent to participate

All procedures performed on animal were approved by the academic committee in Hunan Normal University, Hunan, China (Approval number: 2020E013).

## Conflict of Interests

The authors have declared that no competing interests exist.

## Notes

### Competing Interest Statement

The authors have declared no competing interest.

